# Force sharing between plantarflexor muscles in sheep during treadmill gait

**DOI:** 10.64898/2026.06.23.734066

**Authors:** Stephanie A. Ross, Florian S. Schumacher, Esthevan Machado, Andrew Sawatsky, Timothy R. Leonard, Keiji Hopfner, W. Michael Scott, Fransiska M. Bossuyt, William R. Taylor, Walter Herzog

## Abstract

Muscle force sharing during locomotion is influenced by the mechanical demands of movement and the contractile properties of synergistic muscles. In cats, plantarflexor muscles exhibit distinct functional specialization, with the slow-fibred soleus maintaining relatively constant force across conditions while faster muscles such as the plantaris and gastrocnemius increase force production with increasing locomotor demand. However, it remains unclear whether similar force-sharing patterns occur in larger animals with different musculoskeletal designs. Therefore, the purpose of this study was to examine force sharing between the superficial digital flexor (SDF) and medial gastrocnemius (MG) muscles during treadmill locomotion in sheep. Tendon buckle force transducers were surgically implanted on the SDF and MG tendons of seven sheep, and in vivo muscle forces were recorded during walking and trotting across different speeds and inclines. Both muscles increased force with increasing speed and incline; however, speed had a substantially greater effect than incline. The SDF consistently produced greater absolute force than the MG across all conditions, whereas the MG exhibited slightly larger relative increases in force with increasing speed. Time to peak force decreased with increasing speed in both muscles, although the SDF reached peak force later in stance than the MG across conditions. In contrast to the distinct specialization observed in cats, neither muscle displayed a relatively condition-independent, soleus-like force contribution. These findings suggest that force sharing in sheep is more distributed across synergistic muscles and may reflect the influence of musculoskeletal design, tendon compliance, and mixed fibre-type composition on muscle function in larger species.

## Introduction

Skeletal muscles generate the forces required to move joints during locomotion. Because multiple muscles often contribute to the same joint moment, there are many possible ways to distribute force across synergistic muscles to produce the same movement. This “muscle redundancy” or “force distribution” problem has traditionally been framed as an underdetermined mechanical system in which the number of unknown muscle forces exceeds the number of mechanical degrees of freedom (Crowninshield and Brand, 1981). However, muscles crossing the same joint are not mechanically equivalent. Differences in muscle fibre-type composition, architecture, tendon compliance, and moment arms influence both the mechanical output and energetic cost associated with force production. Consequently, force sharing across muscles is likely constrained by the mechanical capacities and contractile properties of the muscles available to perform a given task (Herzog, 1996).

Muscle fibre-type composition appears to play an important role in determining how forces are distributed across synergists during locomotion. Slow muscle fibres are metabolically and economical well suited for sustained, lower-velocity contractions, whereas faster fibres can generate greater force and power during rapid movements but at a higher energetic cost (Barclay et al., 1993; Crow and Kushmerick, 1982). As contraction velocity and force demands increase, recruitment strategies shift toward faster motor units because slower fibres become less capable of producing force under rapid contractile conditions while still incurring metabolic cost (Hodson-Tole and Wakeling, 2009). Accordingly, muscles with different fibre-type compositions may contribute preferentially under different locomotor demands. Experimental studies in cats provide strong evidence for such functional specialization among plantarflexors. During gait, the soleus, which is composed predominantly of slow fibres, maintains a relatively consistent force contribution across locomotor conditions, whereas the plantaris and gastrocnemius muscles, which contain larger proportions of fast fibres, increase their force output as speed and mechanical demand rise (Herzog et al., 1992; Herzog et al., 1993a; Hodgson, 1983; Prilutsky et al., 1994; Smith et al., 1977; Walmsley et al., 1978). At the highest movement speeds, such as during jumping or paw shaking, activation of the soleus may even decline while the faster muscles dominate force production (Smith et al., 1980; Walmsley et al., 1978). Together, these findings suggest that force sharing across synergistic muscles reflects both the mechanical demands of locomotion and the contractile capacities of the muscles involved.

However, it remains unclear to what extent these patterns of functional specialization generalize across species with different musculoskeletal designs and locomotor demands. Compared to smaller cursorial species such as cats, larger mammals often locomote with more upright limb postures and narrower ranges of joint motion (Biewener, 1989), and possess higher capacity for tendon elastic energy storage (Pollock and Shadwick, 1994). Long compliant tendons can decouple muscle behaviour from whole muscle-tendon unit length changes by allowing elastic stretch and recoil to accommodate much of the joint movement (Biewener et al., 1998; Fukunaga et al., 2001; Hoffer et al., 1989). Similarly, pennate muscle architecture can uncouple fascicle shortening from whole-muscle shortening through fibre rotation and architectural gearing (Eng et al., 2018; Roberts et al., 2019). Together, these features may reduce variability in fibre operating lengths and velocities across locomotor tasks, potentially diminishing the need for highly specialized force-sharing roles among synergistic muscles. Species differences in fibre-type distributions may further alter how muscles contribute across conditions. In contrast to cats, which exhibit strongly polarized fibre-type distributions across plantarflexors, many larger species possess more mixed fibre-type compositions across synergistic muscles (Queeno et al., 2023). Consequently, force-sharing strategies likely vary across species due to differences in locomotor demands, body size, limb geometry, tendon properties, muscle architecture, and fibre-type composition, although the extent to which these factors influence force sharing remains unclear.

Sheep provide an informative model for examining these questions. Sheep are moderate-sized ungulates that typically locomote at slow to moderate speeds and possess relatively upright limb posture and long distal tendons. In addition, sheep lack a substantial soleus muscle, and the superficial digital flexor, which contributes to ankle extension and performs a role analogous to the plantaris in other species, possesses a relatively mixed fibre-type composition compared to the more strongly specialized plantarflexors of cats (Ariano et al., 1973; Konno and Watanabe, 2012). These features suggest that force sharing across the sheep plantarflexors may be less functionally specialized than in species with more distinct fibre-type segregation and locomotor demands. Therefore, the purpose of this study was to examine force sharing between the medial gastrocnemius and superficial digital flexor muscles during treadmill gait in sheep across varying speeds and inclines. Based on the musculoskeletal design and locomotor characteristics of sheep, we hypothesized that neither muscle would exhibit the relatively condition-independent, soleus-like force contribution previously observed in cats. Instead, we predicted that both muscles would increase force production with increasing locomotor demand, reflecting a more distributed and less specialized force-sharing strategy across synergistic plantarflexors.

## Methods

### Animals

We conducted *in vivo* testing on seven healthy female yearling sheep (breed: Île de France; source: commercial producer; weight (mean ± s.d.): 72.5 ± 10.1 kg). Prior to testing, sheep were trained to walk and trot on a motor-driven treadmill for a minimum of 3 months, 3-4 sessions per week, for 10-15 minutes per session. Sheep were housed in individual pens with an open-rail design that allowed for visual and nose contact between sheep in neighbouring pens and were given access to an outdoor paddock for enrichment when weather conditions allowed. The animals had access to water ad libitum and were fed with measured amounts of hay, commercial sheep supplement, and loose minerals. Husbandry and veterinary care were provided and overseen by the University of Calgary Animal Care Unit. All procedures were approved by the University of Calgary Animal Care Committee (protocol AC19-0098).

### Surgery and sensor implantation

The sheep were fasted for 12-15 hours and sedated (0.015 mg/kg Dexdomitor and 2 mg/kg alfaxalone, intravenous) the morning prior to surgery. Animals were deeply anaesthetized with 1- 2% inhaled isoflurane in 1% oxygen, with additional sedation (0.004 mg/kg Dexdomitor, intravenous) as needed. We monitored vital signs throughout surgery. To prep the surgical site, the left hindlimb and back were shaved and cleaned with Hibitane soap and 70% ethanol. We used aseptic technique to reduce the risk of infection throughout the surgery. We made an incision on the lateral aspect of the left hindlimb and used blunt dissection to isolate the medial gastrocnemius (MG; *gastrocnemius, caput mediale*) and superficial digital flexor (SDF; *flexor digitorum superficialis*) muscles and tendons. In sheep and other ruminants, the plantaris is absent or reduced, and the muscle that runs along the posterior lower leg and contributes to ankle extension is the SDF, which has a similar functional role to the plantaris in humans and other vertebrate species. To estimate the muscle forces, we placed an E-shaped buckle force transducer (Herzog et al., 1993b; Walmsley et al., 1978) on each tendon and secured them using nylon monofilament suture. We routed the sensor wires subcutaneously from the hindlimb to the lower spine and soldered them to a custom-made connector with built-in amplifier (AD627, Analog Devices, Wilmington, Massachusetts, USA). Thirty minutes prior to the end of surgery, we discontinued isoflurane and administered intravenous atipamazole (5x volume of Dexdomitor) to reverse the anaesthesia. We also administered analgesics (0.05 mg/kg subcutaneous meloxicam and 0.01 mg/kg intravenous buprenorphine) to reduce post-operative pain. Animals were monitored post-operatively, and analgesics (0.5 mg/kg meloxicam and 0.02 mg/kg buprenorphine, subcutaneous) were provided every 12 hours until the morning of the collection session.

### In vivo data collection

Three to four days following surgery, we collected *in vivo* data during treadmill gait at speeds of 0.67, 1.34, and 1.97 m/s (1.5, 3.0, and 4.4 miles per hour) and inclines of 0, 3, and 6 degrees. To determine the timing of the swing and stance phases of the gait cycle, we video recorded a side view of the treadmill trials at 60 frames per second (NorPix Inc., Montreal, Quebec, Canada).

We recorded the force traces and a sync pulse that was displayed in the videos via a light visible in-frame at 1040 Hz (WinDaq data acquisition software and DI-400 USB DAQ, Dataq Instruments, Akron, Ohio, USA). Once the trials were completed, we sedated the animals (0.015 mg/kg intravenous Dexdomitor) and euthanized them with an overdose of sodium pentobarbital (100 mg/kg intravenous). Following euthanasia, we dissected and isolated the MG and SDF muscles and tendons and collected data to calibrate the force transducers by hanging weights from the ends of the tendons while recording the output voltage.

### Data analysis

We manually tracked the videos to identify the frames that corresponded with the start of the stance and swing phases of the gait cycle and determined the nearest corresponding samples in the force traces. To analyse the force data, we selected 10 consecutive step cycles in the middle of each trial that corresponded to steady-state gait. Due to potential drift in the force transducers between and across trials, we zeroed the output voltages from the sensors at the time corresponding to the lowest voltage per gait cycle. We converted the resultant voltages to force in Newtons using calibration factors obtained from linear calibration curves (force = calibration factor*voltage) for each tendon that we constructed from the calibration trials. To calculate the mean time-varying force trace across the 10 consecutive step cycles, each cycle was first interpolated and resampled to 100 evenly spaced points. The mean force was then calculated at each corresponding point index across the 10 cycles, ensuring that values were aligned by relative time. We then applied this procedure to derive the mean time-varying force for each condition across all sheep. To calculate the mean peak force and time to peak force, we identified the peak force and corresponding time from the start of the cycle for each cycle, and then averaged these values across consecutive steps. Data processing was conducted in MATLAB (MathWorks Inc., Natick, Massachusetts, USA).

To examine differences in maximum force and time to maximum force between conditions and muscles, we used linear mixed-effects models with speed, incline, and muscle as categorical fixed effects, sheep as a random effect to account for repeated measures, and maximum force or time to maximum force as continuous response variables. Type II Wald χ^2^ tests were used to assess the significance of fixed effects. Analyses were conducted in R (version 4.3.2; R Project for Statistical Computing), and models were fit by maximum likelihood using the *lmer* function in the *lme4* package (Bates et al., 2015). Post hoc tests of fixed effects were performed using estimated marginal means with Holm-adjusted p-values (Holm, 1979), implemented via the *emmeans* package (Lenth, 2016; Lenth and Piaskowski, 2026), with simple effects examined for significant interactions. Two animals had partial force data due to issues with the force buckles: one SDF buckle had an insufficient force range, resulting in clipping at high speed and incline conditions, and the wire connection to the MG buckle of another sheep was damaged during the recovery period and did not record forces for any condition. These cases were handled via listwise deletion, as implemented by default in *lme4*.

## Results

Both the MG and SDF forces increased early in stance and then decreased before the foot lifted for swing. While this general pattern of time-varying forces was consistent across conditions for both muscles, there was some step-to-step variability in force magnitude and timing across the ten steps within each trial (Fig. 1). The magnitude of time-varying force across the stance phase was generally higher in both muscles with greater speed (Fig. 2) and incline (Fig. 3). While there were differences between muscle force contributions between the SDF and MG across conditions, as described below, both muscles generally showed increases in force with greater speed and incline (Fig. 4), similar to the cat plantaris and gastrocnemius muscles.

**Figure 1.**
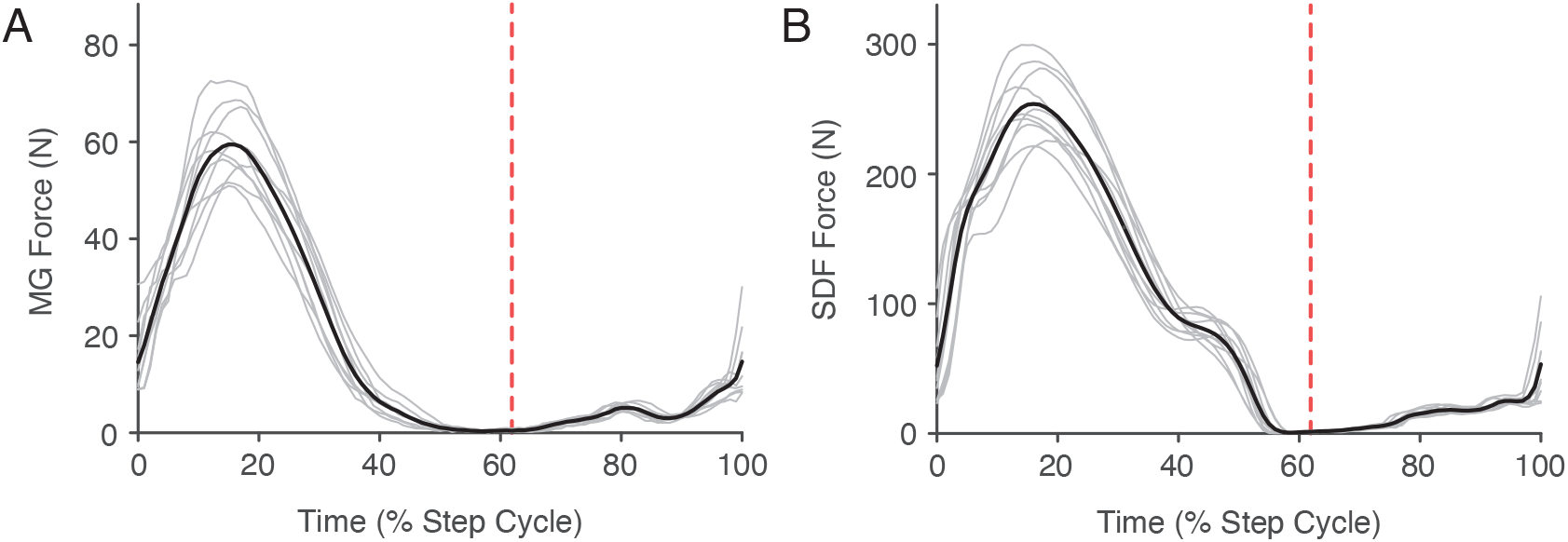
Time-varying muscle force across consecutive step cycles. MG (A) and SDF (B) force over time normalized to the step duration across 10 consecutive step cycles (overlayed in grey) for a representative sheep at 1.34 m/s and 0 degree incline. The mean force across the 10 steps is shown in black. The start of stance occurs at 0% of the step cycle and continues until the vertical dashed red line.

**Figure 2.**
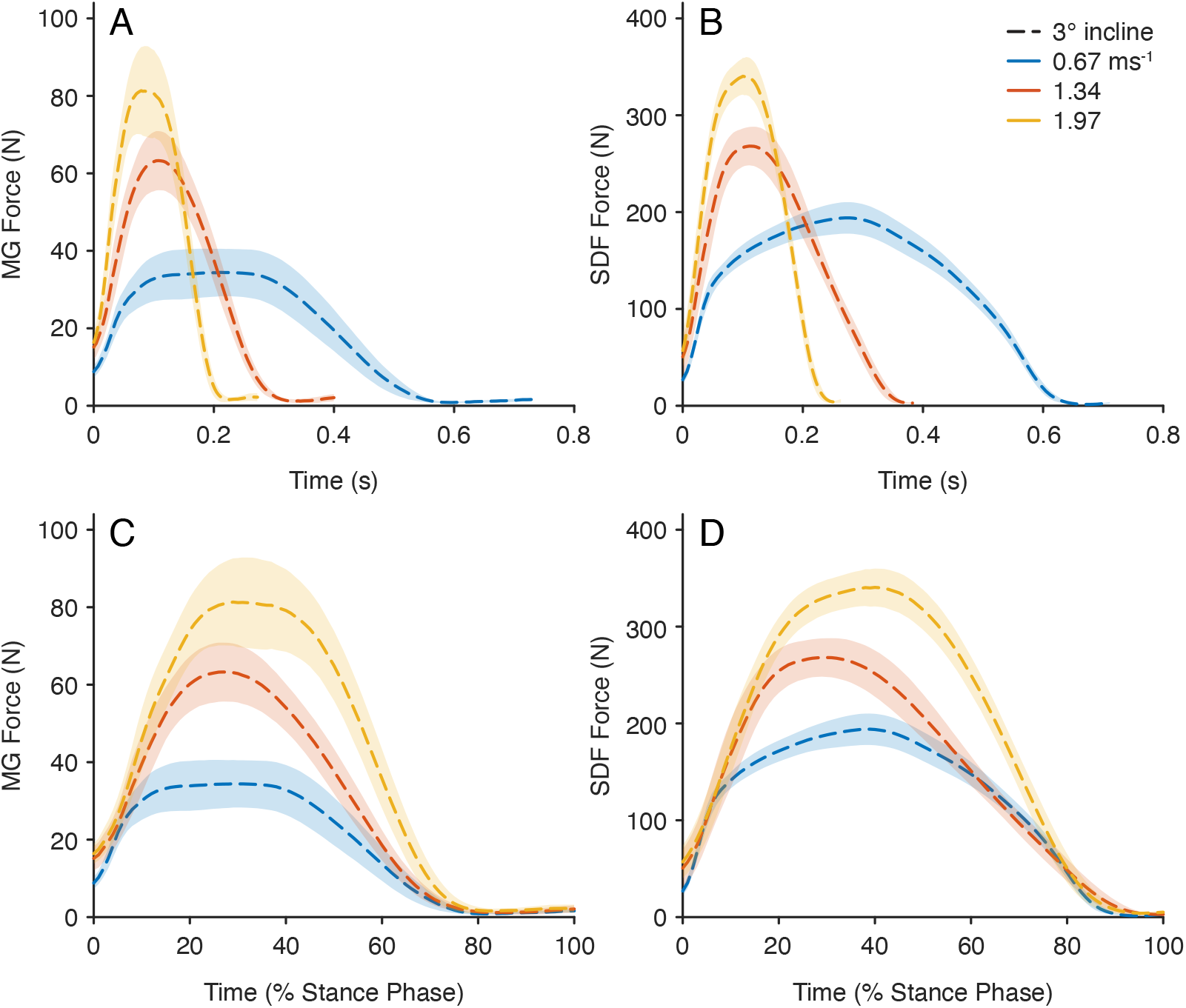
Muscle forces during stance across speeds. Mean MG (A,B) and SDF (C,D) force for all sheep during stance for each speed across absolute time (A,C) and time expressed as a percent of the stance phase (B,D) at 3 degrees incline. Standard errors are shown with the shaded bands. Data are shown from six animals for the MG at all speeds, and from six, five, and five animals for the SDF at 0.67, 1.34, and 1.97 m/s, respectively.

**Figure 3.**
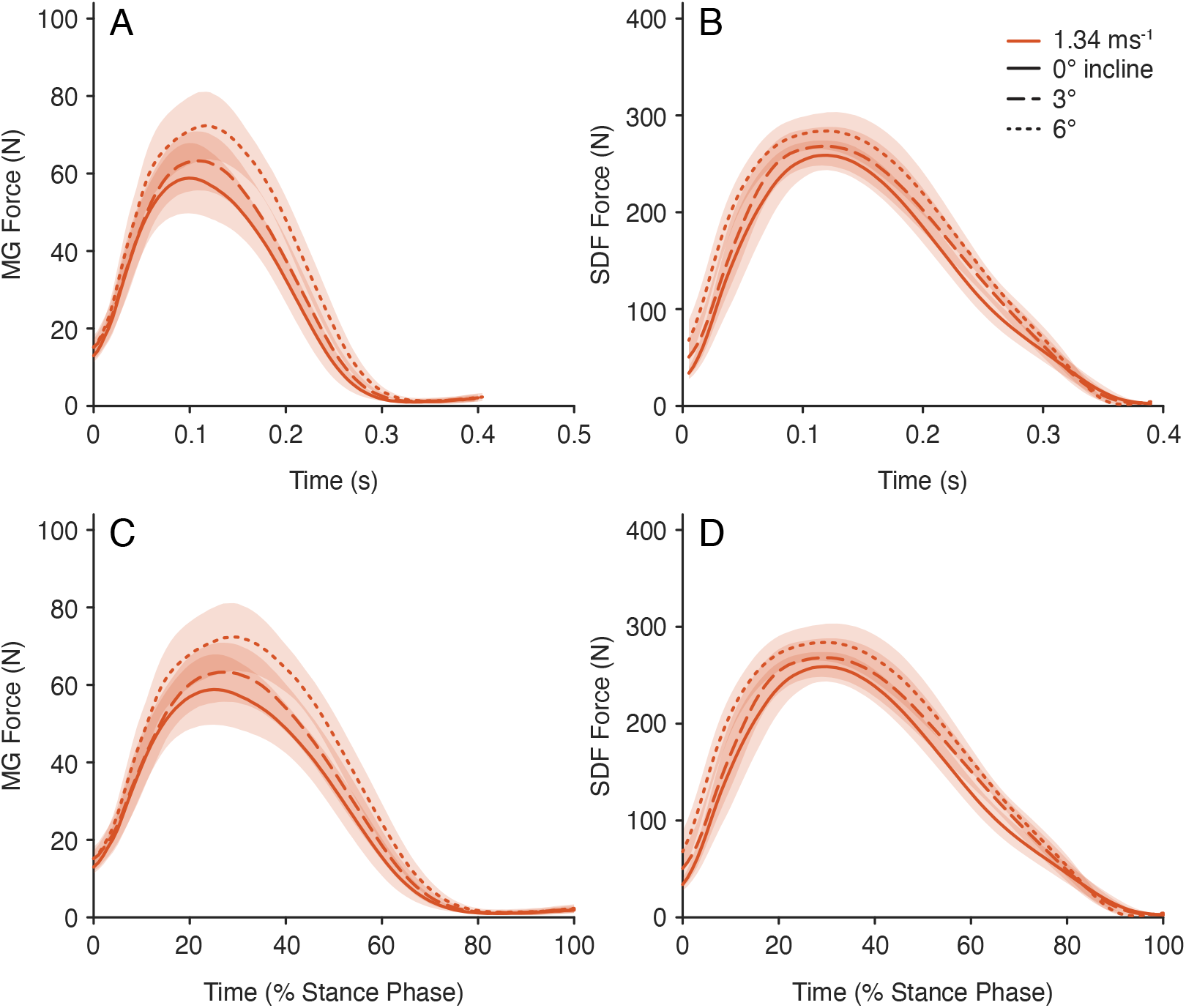
Muscle force during stance across inclines. Mean MG (A,B) and SDF (C,D) force for all sheep during stance for each incline across absolute time (A,C) and time expressed as a percent of the stance phase (B,D) at 1.34 m/s. Standard errors are shown with the shaded bands. Data are shown for six animals for the MG and five for the SDF for all inclines.

**Figure 4.**
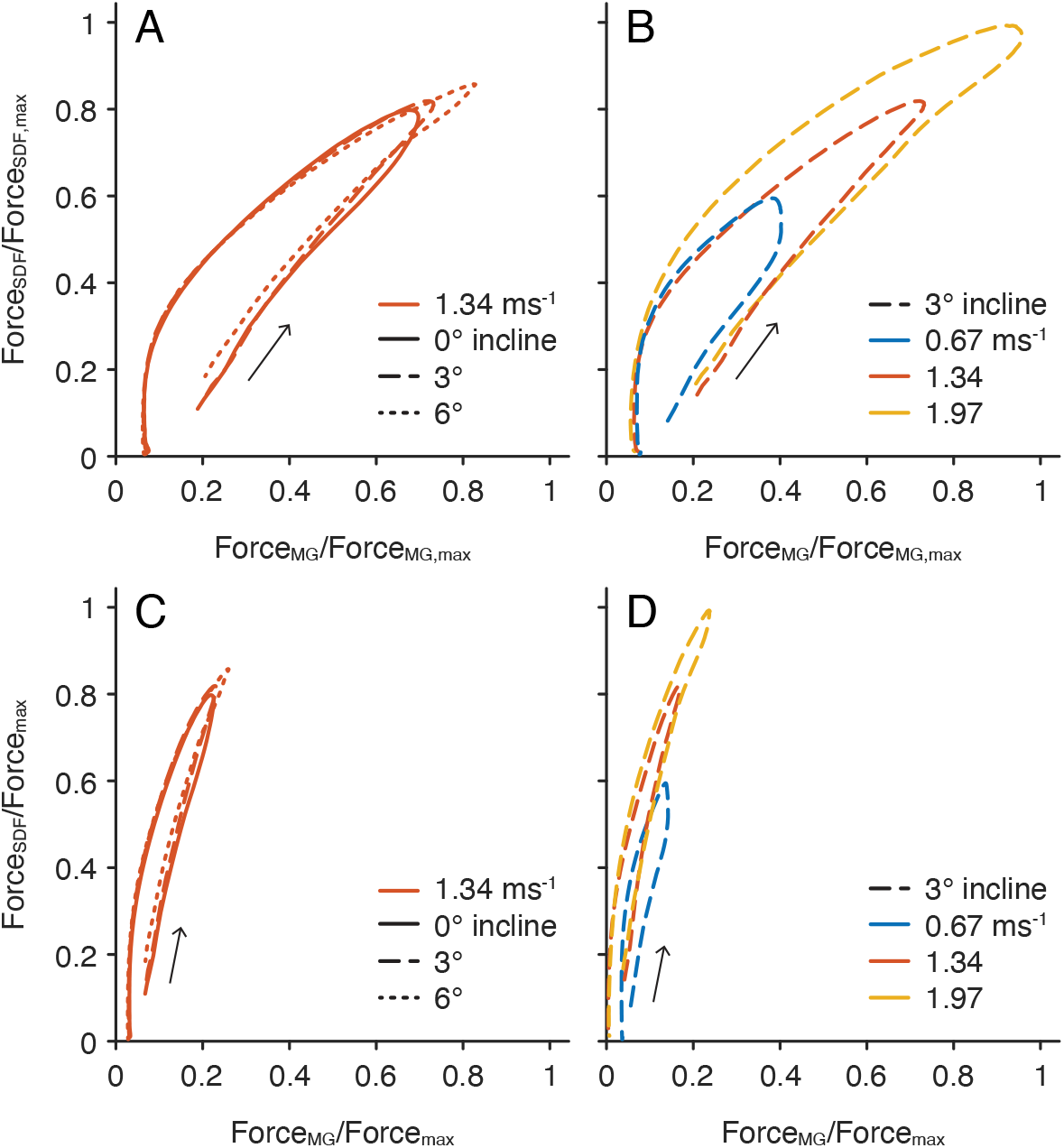
Relationship between SDF and MG force. Mean SDF force versus MG force for all sheep across inclines at 1.34 m/s (A,C) and across speeds at 3 degrees incline (B,D). Forces are shown relative to each muscle’s maximum force across conditions in A and B, and relative to the maximum force for both muscles (SDF in all cases) in C and D.

In terms of specific differences in absolute force (Fig. 5A), results from the linear mixed model showed that force differed significantly between muscles, across speeds, and with incline (muscle: p < 0.001; speed: p < 0.001; incline: p < 0.001). There was a significant interaction between muscle and speed (p < 0.001), but no significant interactions involving incline (muscle × incline: p = 0.77; speed × incline: p = 0.96; muscle × speed × incline: p = 0.99). Linear mixed model estimates confirmed that the SDF muscle produced substantially higher forces than MG across all speeds and inclines (all pairwise contrasts p < 0.001). Peak force increased with speed for both muscles, with post-hoc comparisons showing significant differences between lower (0.67 m/s) and higher speeds (1.34 m/s and 1.97 m/s), particularly for the SDF (e.g., 0.67 m/s vs. 1.97 m/s, SDF: −157.9 ± 14.5 N, p < 0.0001; MG: −53.0 ± 13.8 N, p = 0.0006). In contrast, although incline showed a significant main effect in the model, post hoc tests revealed no significant pairwise differences between incline levels within any muscle × speed combination (all p ≥ 0.22), indicating that incline had minimal practical impact on peak force production.

**Figure 5.**
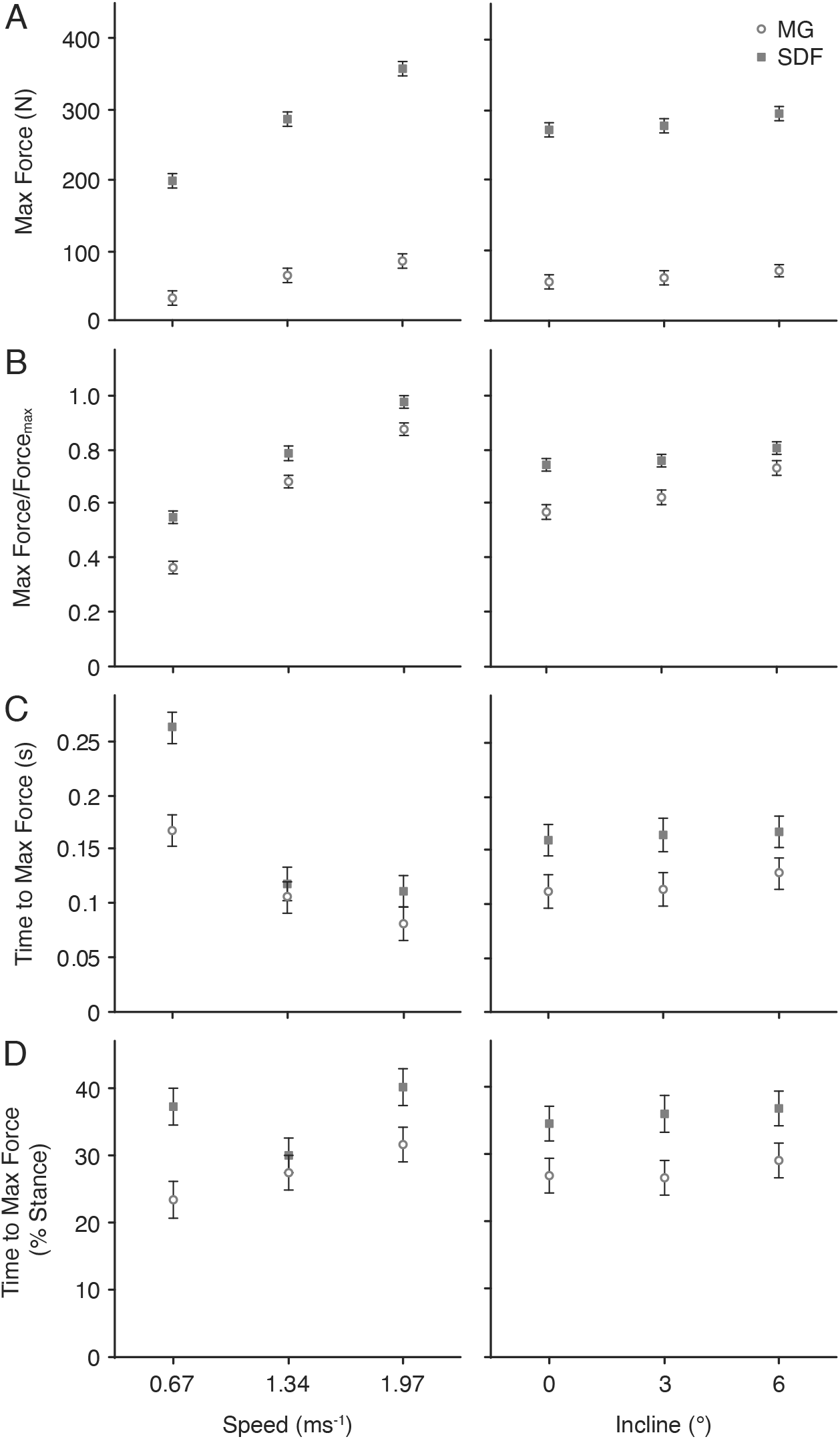
Peak force and timing across speeds, inclines and muscles. Least squares mean maximum absolute force (A), maximum force for each muscle per trial relative to its own maximum across conditions (B), absolute time to maximum force (C), and time to maximum force as a percentage of stance (D) across speeds (left plots) and inclines (right plots) for the SDF (grey squares) and MG (hollow circles). Error bars show the standard error of the mean.

Relative maximum force, normalized to each muscle’s peak across conditions (Fig. 5B), was significantly influenced by muscle, speed, and incline, with significant interactions between muscle × speed and muscle × incline (muscle, p < 0.001; speed, p < 0.001; incline, p < 0.001; muscle × speed, p = 0.014; muscle × incline, p = 0.005). Linear mixed model estimates confirmed that the SDF generally achieved higher relative peak forces than the MG, particularly at lower inclines and speeds (e.g., 0.67 m/s, incline 0: −0.24 ± 0.04, p < 0.0001), while at the steepest incline (6°), differences diminished and were non-significant at higher speeds (speed 1.34 m/s and 1.97 m/s, p > 0.22). Contraction speed strongly increased relative peak force in both muscles, with significant pairwise differences across all speed levels (e.g., MG, 0° incline: 0.67 m/s vs. 1.97 m/s = −0.51 ± 0.04, p < 0.001; SDF, 0° incline: 0.67 vs. 1.97 m/s = −0.41 ± 0.05, p < 0.001). Incline effects were muscle-dependent: in the MG, relative maximum force increased significantly at the steepest incline across all speeds (e.g., 1.97 m/s: 0° incline – 6° incline = −0.189 ± 0.041, p < 0.001), whereas the SDF showed no significant incline-related changes (all p > 0.28). Overall, peak force output was primarily determined by muscle and speed, with incline modulating force mainly in the MG.

Time to peak force, expressed both in absolute terms (Fig. 5C) and relative to stance duration (Fig. 5D), differed significantly between muscles and across speeds, but was minimally affected by incline. We found significant main effects of muscle (absolute: p < 0.001; relative to stance: p < 0.001) and speed (absolute: p < 0.001; relative: p < 0.001), and a significant muscle × speed interaction (absolute: p < 0.001; relative: p < 0.001), whereas effects of incline and interactions involving incline were not significant. Post hoc contrasts indicated that the SDF reached peak force later than the MG across most speed and incline conditions. For example, at 0° incline and 0.67 m/s, the MG peaked at 0.15 ± 0.02 s (21.1 ± 3.0% stance) while the SDF peaked at 0.26 ± 0.02 s (36.3 ± 2.7% stance, p < 0.001). Increasing speed consistently decreased absolute time to peak force in both muscles (e.g., MG: 0.151 s to 0.215 s; SDF: 0.258 s to 0.407 s), but because stance duration decreased at faster speeds, time to peak force as a percentage of stance increased (e.g., MG: 21.1% to 32.7%; SDF: 36.3% to 40.8% of stance at 0° incline). The effects of incline were minor; only the MG at 6° showed a slight increase in absolute time (0.151 to 0.260 s, p = 0.038), with no significant changes for the SDF. In summary, muscle and speed were the primary determinants of peak force timing, with the MG reaching peak force earlier than the SDF both in absolute time and relative to stance, while incline had minimal influence.

## Discussion

The purpose of this study was to examine force sharing in the SDF (homologous to plantaris) and MG muscles of sheep during treadmill gait across varying speeds and inclines. Previous research in cats has shown that while plantaris and gastrocnemius muscles both increase force contribution with greater speeds and inclines (Fig. 6B), this is not the case for the soleus, which largely maintains its force contribution across speeds (Herzog et al., 1993a). In sheep, where the soleus is negligibly small (Konno and Watanabe, 2012), we hypothesized that the MG and SDF would have similar force behaviour as in cats, such that their forces would both proportionally increase with greater mechanical demand of gait. Consistent with our hypothesis, we found that while there were significant differences in how to the two muscles alter their force in response to increases in gait speed and incline, both muscles in the sheep showed generally greater forces with greater speed and incline (Fig. 4). An alternative possibility was that one of these muscles might adopt a more soleus-like role, maintaining a relatively constant force contribution across conditions. Our findings do not support this alternative, as neither muscle demonstrated a consistent, condition-independent force output.

**Figure 6.**
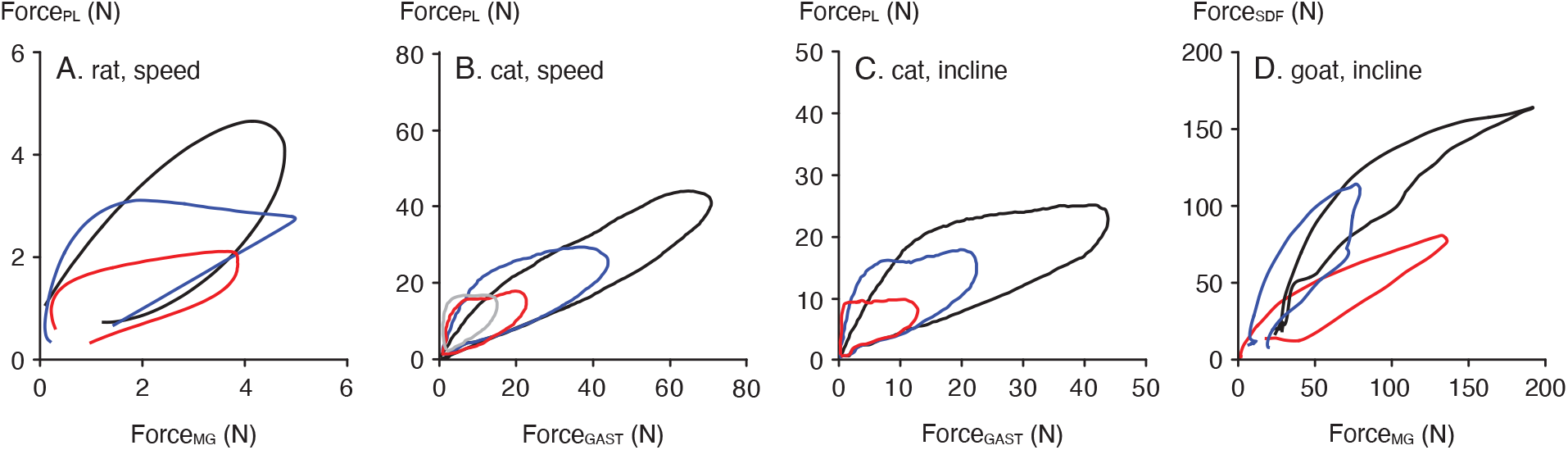
Comparison of force sharing across species. (A) rat plantaris versus MG forces across speeds (red = walk, blue = trot, black = gallop; Eng et al., 2019; permission pending); (B) cat plantaris versus gastrocnemius forces across speeds (grey = 0.4 m/s, red = 0.7 m/s, blue = 1.2 m/s, black = 1.8 m/s; Herzog et al., 1993a; permission pending); (C) cat plantaris versus gastrocnemius forces across inclines at 0.7 m/s (red = 10 degrees decline, blue = level, black = 10 degrees incline; Herzog et al., 1993a; permission pending); (D) goat SDF versus MG forces across inclines at a trot of 2.5 m/s (red = 15 degrees decline, blue = level, black = 15 degrees incline; McGuigan et al., 2009; permission pending). Each subplot traces shows data from one representative animal.

The objective function that governs how force is distributed across muscles likely incorporates muscle-specific properties relative to the contractile conditions, including fibre-type composition. The soleus is predominantly composed of slow fibres, whereas the plantaris is primarily fast-fibred and the medial gastrocnemius is either mixed or fast-fibred in most species (Ariano et al., 1973; Edgerton et al., 1975; Peter et al., 1972; Queeno et al., 2023). According to the size principle (Henneman et al., 1965a; Henneman et al., 1965b), smaller motor units consisting of slower fibres are recruited before larger motor units composed of faster fibres. This pattern of motor unit recruitment within muscles appears to be paralleled by recruitment of muscles within synergistic groups (Wakeling et al., 2011). Consequently, muscles rich in slow fibres tend to contribute preferentially when overall force demands are low, while increasing force requirements necessitate the recruitment of faster motor units. This pattern can shift for higher contraction speeds, such that faster motor units are preferentially recruited first because slower motor units are less, or not, capable of producing force under rapid conditions yet still incur a metabolic cost (Hodgson, 1983; Hodson-Tole and Wakeling, 2009; Wakeling et al., 2006; Wakeling et al., 2012). These patterns align with findings from previous work in cats, where the soleus, composed entirely of type I (slow) fibres, is activated at slower speeds, while the plantaris and gastrocnemius, which have a higher proportion of fast fibres (Ariano et al., 1973), become increasingly active with increasing speed as force demands rise (Herzog et al., 1992; Herzog et al., 1993a; Hodgson, 1983; Prilutsky et al., 1994; Smith et al., 1977; Walmsley et al., 1978). At the fastest gait speeds, there can be a decline or inhibition of soleus activation, an effect that is most pronounced at the fastest speeds that occur in movements such as jumping (Walmsley et al., 1978) and paw shaking (Smith et al., 1980). Overall, these observations may suggest that muscle recruitment strategies are influenced by fibre-type composition and contractile demands, with a shift toward faster motor units as both force and speed requirements increase.

In contrast to cats, which exhibit strongly polarized fibre-type distributions across muscles, the sheep SDF and MG both contain mixed fibre types, with the SDF having a relatively greater proportion of slow fibres compared to the MG (Konno and Watanabe, 2012). Although both muscles increased force production as speed increased, and the SDF generated greater absolute force than the MG under all conditions, the MG showed slightly larger relative increases in force with increasing speed (Fig. 5). This pattern may reflect the comparatively slower fibre-type composition of the SDF, causing it to behave somewhat like the soleus in cats by maintaining more consistent force output across speeds, whereas the faster-fibred MG is better suited to augment force as demand rises. A similar fibre-type-related pattern may also exist among the cat plantarflexors (Fig. 6B), where the slightly faster-fibred gastrocnemius (Ariano et al., 1973) exhibits relatively larger increases in force with increasing speed compared to the plantaris (Herzog et al., 1993a). In contrast, studies in rats have reported relatively larger speed-related increases in force in the plantaris compared to the MG despite similar fibre-type compositions between the two muscles (Fig. 6A; Eng et al., 2019). This seemingly weaker relationship between fibre-type composition and force contribution across speeds in rats may indicate that other factors, such as MTU properties or differences in contractile conditions between muscles, play a larger role in determining force sharing patterns in this species compared to cats and sheep.

Differences in the timing of force production across speeds in the sheep further support the idea that the SDF functioned more like a slower muscle than the MG. The absolute time to peak force decreased for both muscles at higher speeds; however, the reduction became smaller for the SDF, such that the difference in time to peak force between the intermediate and highest speeds was minimal (Fig. 5C, left). This again may be attributable to the slower fibre composition of the SDF, as slow fibres generally take longer to reach peak activation and force following excitation (Burke et al., 1973); thus, they may have reached the limit of how early they can attain peak force relative to the start of stance without excessive preactivation. Notably, this effect was not observed with increases in incline (Fig 5C, right), suggesting that this constraint is more specific to speed-related demands rather than force requirements. If motor unit recruitment were examined, a pattern more similar to that observed in cats at the muscle level might emerge at the motor unit level, whereby slower fibres are recruited first and remain relatively constant across conditions, while faster motor units increase their activation to modulate force as locomotor demand rises. This increase in recruitment of faster motor units in mixed muscles with increasing speed has been observed previously in other species, including rats (Hodson-Tole and Wakeling, 2008), goats (Lee et al., 2012), and humans (Wakeling et al., 2006). Together, these findings suggest that, in contrast to the distinct fibre-type-driven specialization of muscles in cats, sheep may exhibit a recruitment strategy in which slow soleus-like and fast motor unit behaviours are distributed across muscles rather than segregated between them.

Differences in muscle force sharing across species may reflect variation in contractile demands arising from distinct anatomical and locomotor conditions. For example, sheep have longer limbs and longer distal tendons compared to smaller animals such as cats and rats, which act to decouple muscle fibre behaviour from limb mechanics, as the compliant tendons can accommodate a substantial portion of the MTU length change instead of the muscle fibres themselves (Alexander, 2002). As a result of this uncoupling, the contractile conditions experienced by the muscles with long tendons may remain relatively constant across gait speeds, because changes in limb mechanics are largely taken up by the tendon (Biewener, 1998; Biewener et al., 1998; Harrison et al., 2010; Lai et al., 2014). Larger species such as sheep also operate with a more extended limb posture and a narrower range of joint angles (Biewener, 1989), which reduces variability in muscle moment arms and fibre operating lengths across tasks. Moreover, because faster muscle fibres could incur a greater inertial penalty such that their mechanical advantage at faster speeds converges with that of slower fibres as body and muscle size increases (Ross and Wakeling, 2016; Ross and Wakeling, 2021; Ross et al., 2018), this may further diminish the functional benefit of fibre-type specialization in larger animals. Whether a muscle is mono- or biarticular and the orientation of its force vector may also influence force sharing. This has been shown in cats, where the monarticular soleus primarily contributes to vertical support and stabilization whereas the biarticular MG is more involved in controlling the direction of the ground reaction force (Kaya et al., 2006). This partitioning of mechanical roles between muscles may be reduced in sheep, either due to reduced necessity or redistribution across other muscles with different origins and insertions. Together, these factors may reduce the necessity for specialized mechanical roles, such as muscles that maintain relatively constant force across conditions like the cat soleus, in larger species such as sheep.

An additional factor that may contribute to differences in force sharing across species is variation in muscle architecture and its scaling with body size. As animals increase in size, muscles tend to become relatively shorter and more pennate, with shorter fascicles (Bishop et al., 2021; McGowan et al., 2008) and higher pennation angles (Dick and Clemente, 2016) compared to those in smaller species, although evidence for these trends comes from a limited number of species and broader comparative analyses remain sparse. Pennate architecture can uncouple fibre (or fascicle) shortening from whole-muscle shortening, such that changes in muscle length can be accommodated by fibre rotation rather than fibre shortening (Eng et al., 2018; Roberts et al., 2019). As a result, fibres can shorten more slowly than the whole muscle, allowing them to operate at more favourable regions of the force-velocity curve and potentially increasing force output. Although this is offset by reduced force transmission along the line of action, since only the component of the fibre force aligned with the muscle longitudinal axis is transmitted and this decreases with increasing fibre rotation, the increase in fibre force associated with more optimal force-velocity conditions may outweigh this geometric loss. In the context of the present study, such potential architectural scaling provides a complementary mechanism to tendon-mediated effects in explaining the relatively uniform force sharing patterns observed across muscles in sheep. In particular, the combination of more pennate muscle architecture and reduced coupling of fibre and muscle length changes and velocities may contribute to more constrained fibre operating conditions across locomotor tasks, further reducing the need for muscle fibre-type specialization.

This study has several limitations that should be considered when interpreting the results. First, we did not measure muscle fibre rotations, fibre lengths, or tendon lengths, and therefore our inferences regarding how MTU properties and contractile conditions may dictate differences in force sharing patterns across species are based on prior literature and mechanical reasoning rather than direct measurements. Second, we assumed that the maximum force for each muscle occurred under the experimental condition that produced the highest measured force. However, it is possible that the true peak muscle force was not captured within the tested range of speeds and inclines, meaning that our estimates of relative force-sharing patterns may not fully reflect each muscle’s absolute functional capacity. Third, we did not measure forces from the lateral gastrocnemius, as the lateral gastrocnemius and MG tendons were separated during implantation of the force buckles. Consequently, the role of the lateral gastrocnemius in distributing force across the plantarflexor group remains unclear. Future work could combine measurements of muscle fibre length changes, tendon behaviour, and individual muscle forces to quantify how force contribution is dictated by contractile conditions, and how these relationships may vary across species.

The findings of this study have broader implications for understanding muscle force sharing and apparent redundancy across species. Muscle design and function are not only dictated by animal morphology, such as limb geometry and species-specific locomotor kinematics, but may also be shaped by tendon and muscle properties and behaviour that can uncouple muscle fibre from joint-level mechanical requirements. In the present study, force sharing patterns across the plantarflexor group were more uniform in sheep than those observed previously in cats, suggesting reduced specialization in muscle roles in sheep relative to cats. Accordingly, reductionist approaches that assume multiple muscles crossing a joint can be represented by a single equivalent actuator may not fully reflect the complexity of muscle-tendon interactions that govern force sharing across conditions. However, such simplifications may become more reasonable in larger animals where long tendons and limited kinematic variability reduce the need for highly differentiated muscle roles. More broadly, these results contribute to understanding how musculoskeletal design is shaped by mechanical function, suggesting that differences in muscle properties and force sharing patterns, for example the lack of a distinct soleus in sheep, may reflect reduced mechanical necessity for such specialized roles rather than a loss of function.

## Acknowledgements

We thank the veterinary and husbandry staff at the University of Calgary Veterinary Science Research Station for their assistance with animal care and training. We also thank Hoa Nguyen and Andrzej Stano for their help with data collection.

## Funding

Funding for this study was provided by the Natural Sciences and Engineering Research Council of Canada (NSERC) and the Nigg Chair for Mobility and Longevity to WH. Salary support for F.S.S. was provided by the AO Foundation and the ETH Zurich Foundation. S.A.R. was supported by a Banting Postdoctoral Fellowship and a Canadian Institutes of Health Research (CIHR) Postdoctoral Fellowship.

## Notes

### Competing Interest Statement

The authors have declared no competing interest.

